# Sucrose rather than GA transported by AtSWEET13 and AtSWEET14 supports pollen fitness at late anther development stages

**DOI:** 10.1101/2022.05.05.490840

**Authors:** Jiang Wang, Xueyi Xue, Houqing Zeng, Jiankun Li, Li-Qing Chen

## Abstract

**Summary:**

- Both sugar and hormone gibberellin (GA) are essential for anther-enclosed pollen development and thus for plant productivity in flowering plants. Arabidopsis (*Arabidopsis thaliana*) AtSWEET13 and AtSWEET14, which are expressed in anthers and associated with seed yield, transport both sucrose and GA. However, it is still unclear which substrate transported by them directly affects anther development and seed yield.
- Histochemical staining, cross-sectioning and microscopy imaging techniques were used to investigate and interpret the phenotypes of *AtSWEET13* and *AtSWEET14* double mutant during anther development. Genetic complementation of *atsweet13;14* using AtSWEET9 that transports sucrose but not GA was conducted to test the substrate preference relevant to the biological process.
- The loss of *AtSWEET13* and *AtSWEET14* resulted in reduced pollen viability and therefore decreased pollen germination. AtSWEET9 fully rescued
- the defects in pollen fertility of *atsweet13;14*, indicating AtSWEET13/14 mediated sucrose rather than GA is essential to pollen fertility.
- AtSWEET13 and AtSWEET14 mainly function at the anther wall during late anther development stages and are likely responsible for sucrose efflux into locules to support pollen development to maturation, which is vital for subsequent pollen viability and germination.

## Introduction

Pollen development and formation are vital for the reproduction of flowering plants (Ma, 2005). Developing pollen requires a sugar supply from the somatic cell layers of anthers since immature pollen is well surrounded by locules in the anthers. The somatic cell layers consist of the outermost epidermal cell layer, the endothecium, the middle cell layer, and the innermost tapetum (van der Linde & Walbot, 2019). The male gametophyte is symplasmically isolated from the anther cells, resulting from dissociation between meiotic cells and the tapetum (Ma, 2005; Borghi & Fernie, 2017). Plasma membrane-localized carrier proteins are needed for sugar export from the somatic cell layers of the anther and import into pollen (Borghi & Fernie, 2017). Sucrose is symplasmically unloaded into the connective tissues of the anthers from the phloem, largely following a long-distance sugar transport starting from source tissues. Sucrose is either exported to the locules directly or hydrolyzed into hexoses by an invertase before exported to the locules. However, sugars have to cross different layers in the anther wall to reach the locules. Tapetal cells are structurally disconnected from the adjacent middle cell layer (Clément & Audran, 1995), which indicates sugars must be apoplasmically transported from the middle cell layer into the tapetum cells, and from there, are exported into the locules and finally reach the pollen.

Sugars Will Eventually Be Exported Transporters (SWEETs), primarily transporting both mono- and di-saccharides, are involved in many biological processes associated with apoplasmic transport routes, such as phloem loading and nectar secretion (Chen *et al*., 2012; Lin *et al*., 2014,; Xue *et al*., 2022). Sucrose Uptake Transporters (SUTs/SUCs), mainly transporting sucrose, are often paired with SWEETs in many processes (Braun, 2022). In tomato, SlSWEET5b mediates hexose export into the locules; unsurprisingly, the *SlSWEET5b* silencing mutant exhibits reduced pollen germination and seed production (Ko *et al*., 2022). In rice, OsSUC1 is highly expressed in the wall of anther, and the *ossuc1* mutant shows impaired pollen function without affecting pollen maturation (Hirose *et al*., 2010). OsSWEET11a and OsSWEET11b are expressed in anther veins, and *ossweet11a;11b* mutant is male sterile (Wu *et al*., 2022). In cucumber, the knockdown mutant of tapetum- and pollen-localized *CsSUT1* shows male sterility (Sun *et al*., 2019). AtSUC1 is expressed in the connective tissue of anther and mature pollen, and the *atsuc1* mutant produces defective pollen resulting in a low pollen germination rate (Stadler *et al*., 1999; Sivitz *et al*., 2008). In Arabidopsis, glucose transporter AtSWEET8 (RPG1; Ruptured Pollen Grain 1) and sucrose transporter AtSWEET13 (RPG2) with transcripts detected in the tapetum of anther are involved in primexine deposition, microspore development, and subsequent seed formation (Guan *et al*., 2008; Sun *et al*., 2013). The mutant *rpg1* (*atsweet8*) exhibits defective microspore development, and the double mutant *rpg1rpg2 (atsweet8;13)* results in an almost sterile phenotype (Sun *et al*., 2013). Besides sugars, developing pollen also requires gibberellin (GA) for its viability and development (Plackett *et al*., 2011). The tapetum of the anthers and pollen grains are positioned as major sites for GA synthesis during flower development (Itoh *et al*., 1999). For example, the Arabidopsis GA biosynthesis-deficient mutant *ga1-3* is male sterile (Sun *et al*., 1992). Similarly, the triple mutant *gid1a;b;c* of GA receptor GID1 (Gibberellin-Insensitive Dwarf 1) showed a dwarf phenotype and male sterility (Griffiths *et al*., 2006). GA movement across different cells or tissues has been reported in many biological processes, which often involve GA transporters (Binenbaum *et al*., 2018). Deficiency in GA transport also affects anther development and fertility. For instance, AtGTR1/NPF2.10 (Glucosinolate Transporter 1; Nitrate transporter 1/Peptide transporter Family 2.10) transports jasmonoyl-isoleucine (JA-Ile) and GA in addition to glucosinolates; The *gtr1* mutant has reduced fertility due to impairment in filament elongation and anther dehiscence, and these phenotypes can be rescued by exogenous GA application (Saito *et al*., 2015).

Unexpectedly, plasma membrane-localized AtSWEET13 and AtSWEET14 have been shown to transport GA (Kanno *et al*., 2016) after they were well characterized as sucrose transporters (Chen *et al*., 2012). Besides AtSWEETs, OsSWEET3a has also been found to transport GA in addition to 2-deoxy-glucose (Morii *et al*., 2020). The *atsweet13;14* double mutant has been shown to have a defect in anther dehiscence, which can be rescued by exogenous application of an excess amount of GA (Kanno *et al*., 2016). These results suggested GA might be the predominant substrate of AtSWEET13 and AtSWEET14 in anther. However, it is uncertain whether this rescue is due to altered GA transport or due to an indirect effect of GA on sugar transport because GA and sugar are interconnected in signaling pathways of many developmental processes, including flowering (Matsoukas, 2014). To clarify which substrate of AtSWEET13 and AtSWEET14 is important in anther to support pollen development, we used AtSWEET9, with sucrose transport activity but does not have GA transport activity (Lin *et al*., 2014; Kanno *et al*., 2016; Wu *et al*., 2022), as a genetic tool to complement *atsweet13;14* under control of the AtSWEET14 promoter. As a result, AtSWEET9 was able to fully rescue the defected pollen germination phenotype of *atsweet13;14*. Our data suggest that sucrose transported by AtSWEET13 and AtSWEET14 is vital to pollen viability and male fertility in Arabidopsis. In addition, our cross-section results from the GUS reporter line and starch accumulation comparison between Col-0 and *atsweet13;14* support that *atsweet13;14* mediated sucrose release is from the endothecium into the locule at the late stages of the anther development when the tapetum has degenerated.

## Materials and Methods

### Plant materials and growth conditions

The Arabidopsis *(Arabidopsis thaliana)* Col-0 plants were grown under a controlled condition with a constant temperature of 22°C, the light intensity of 100~150 μmol m^−2^ s^−1,^ and a 16-h light / 8-h dark photoperiod. T-DNA mutants of *atsweet13* (*SALK_087791*), *atsweet14* (*SALK_010224*) were obtained from ABRC. Homozygous lines were genotyped using primers of P1-P4 (Supporting Information **Table S1**) and used in related experiments. The floral dip transformation method (Clough & Bent, 1998) was used to generate all the transgenic lines described in this study. At least 16 T1 lines were generated for each construct, and at least 2 randomly selected lines were propagated to generate T3 homozygous seeds.

### Constructs for localization and complementation

The 5’ upstream promoter fragments before *ATG* of *AtSWEET13* (2083 bp) and *AtSWEET14* (3798 bp) were amplified using gene-specific primers (P5-P8) with an *Xba*I site added to the reverse primers. The promoter fragments each were cloned into the entry vector pDONR221-f1 via the BP reaction to make pDONR221-f1/*pSWEET13* and *pDONR221*-f1/*pSWEET14*. The CDS of *AtSWEET13*, *AtSWEET14*, and *AtSWEET9* each was amplified using their own specific primers (P9-P14) and then seamlessly subcloned to generate the corresponding constructs pDONR221-f1/*pSWEET13:SWEET13*, pDONR221-f1/*pSWEET14:SWEET14*, and pDONR221-f1/*pSWEET14:SWEET9* by In-Fusion^®^ (TaKaRa, Shiga, Japan) after the linearization at the *XbaI* restriction site. The resulting entry vectors were subjected to LR reaction with the destination vectors pBGGUS and pHGY (Kubo *et al*., 2005) to make *pBGGUS/pSWEET13:SWEET13-GUS, pBGGUS/pSWEET14:SWEET14-GUS*, pHGY/*pSWEET13:SWEET13-YFP*, pHGY/*pSWEET14:SWEET14-YFP*, pHGY/*pSWEET14:SWEET9-YFP*. For the GUS localization assay, pBGGUS constructs were transformed into Col-0. For complementation and fluorescence assay, pHYG constructs were transformed into *atsweet13;14* double mutant.

### Reciprocal crossing and silique imaging

Pollen from either Col-0 or *atsweet13;14* was taken to cross over with the stigma of either Col-0 or *atsweet13;14*. Siliques were cleared with a solution containing 1% SDS and 0.2 M NaOH for 2 days followed by imaging using a dissecting microscope after they were collected 14 days from crossing. At least 5 siliques were generated from each combination for each of the three independent trials. Two representative siliques for each combination were shown.

### *In vitro* pollen germination

The *in vitro* pollen germination assay was conducted according to a previously described method (Wang *et al*., 2022b). Three independent repeats were conducted.

### Soluble sugar quantification

Pollen grains from 100 flowers at stage 13 were used for soluble sugar quantification. Each sample was treated with 1 ml ethanol (80% v/v) and then mixed by vortexing for 30 s. The samples were kept at 80°C for 40 min and inverted to mix every 10 min. The supernatant was transferred into a new tube after being centrifuged at 16,000 × g for 15 min. The extraction process was repeated twice for each sample. The sugar-containing supernatant was kept 42°C for evaporation before re-dissolved in 0.15 ml milli-Q water. After infiltration using a nylon membrane (0.22 μm), the samples were analyzed by high-performance liquid chromatography (HPLC) (Agilent Technologies, 1200 series, Santa Clara, CA) using a separation column containing Rezex™ RCM-Monosaccharide Ca^2+^ (8%) (Phenomenex, Torrance, CA). The average of three independent repeats was used for calculation.

### Fatty acids measurement

The total fatty acid profiles from ~0.5 mg freeze-dried pollen samples were determined using a direct acid-catalyzed transmethylation method with minor modifications (Browse *et al*., 1986). Samples were placed in Teflon-lined screw-capped glass tubes containing 1 ml 5% H2SO4 (v/v) in methanol (freshly prepared), 50 μg butylated hydroxytoluene (BHT), and 5 μg C15:0 pentadecanoic acid as internal standard, and then mixed with 300 μl of toluene as cosolvent followed by vortexing for 60 s. Samples were heated at 85°C for 2h. After cooling down to room temperature, 1 ml 0.9% NaCl (w/v) and 1 ml hexane were used to extract fatty acid methyl esters (FAME). Samples were vortexed for 60 s and centrifuged to accelerate phase separation. The upper organic phase (FAME containing) was transferred into a 2 ml injection vial. Samples were injected into a GC (Agilent, Santa Clara, CA) with a flame ionization detector (FID) using DB-Wax 30 m column (30 m × 0.25 mm × 0.25 um; J&W Scientific, Folsom, CA). The GC conditions were set as follows: split mode injection (1:2); injector and flame ionization detector temperature, 260°C; oven temperature program 150°C for 3 min, then increasing at 10°C min^−1^ to 240°C and holding this temperature for 5 min. The average of four independent repeats was calculated.

### GUS histochemical analysis

GUS staining was performed as previously described (Chen *et al*., 2012). Entire inflorescences of 6-week-old plants were collected for histochemical GUS staining.

### Paraffin-embedded cross-sectioning

Paraffin-embedded cross-sectioning was performed as described with minor modifications (Zhang *et al*., 2021). GUS-stained samples of entire inflorescence from *pSWEET13:SWEET13-GUS* and *pSWEET14:SWEET14-GUS* plants were washed three times by PBS (pH 7.2) and fixed with cold 4% paraformaldehyde and 2.5 % sucrose buffer overnight. Samples were then embedded in paraffin, followed by sectioning and imaging as previously described (Zhang *et al*., 2021).

### Microscopy imaging

A Zeiss Apotome.2 (Carl Zeiss, Thornwood, NY, USA) was used for fluorescent signal acquisition. Due to strong autofluorescence observed under the YFP filter set, the GFP filter set was used to lower the autofluorescence background. Different FL filter sets (GFP: 470/20 nm excitation, 505–530 nm emission, RFP: 546 nm excitation-590 nm emission) were used to image samples as needed. Image acquisition parameters were held consistent. For the confocal microscope, all images were taken using LSM 710 (Carl Zeiss, Thornwood, NY, USA). Argon laser excitation wavelength and emission bandwidths were 514 nm (10% intensity) and 518–565 nm for YFP (gain 750), 514 nm (10% intensity) and 619–697 nm for chlorophyll autofluorescence (gain 450), respectively. Field-Emission Environmental Scanning Electron Microscope (ESEM-FEI; Thermo Fisher Scientific, MA, USA) with Energy-Dispersive Spectroscopy (EDS) on Wet mode was used for SEM imaging of anthers at different stages.

### Pollen viability assay

Pollen viability assay was conducted as previously described (Muhlemann *et al*., 2018) with minor modifications. For each repeat, 50 flowers at stage 13 for each genotype were collected into a 2 ml tube, immersed in 1 ml of liquid pollen viability solution (PVS, 290 mM sucrose, 1.27 mM Ca(NO_3_)_2_, 0.16 mM boric acid, 1 mM KNO_3_) and vortexed vigorously for 60 s to release mature pollen grains into the solution. Pollen from each genotype were collected after being centrifuged at 15,000 g for 1 min. The pollen pellet was resuspended in 1 ml PVS containing 0.001% (w/v) fluorescein diacetate (FDA), and 10 μM propidium iodide (PI). Pollen was stained for 15 min at 28°C and then centrifuged. The pollen pellet was washed using PVS followed by imaging using a fluorescence microscope. FDA was imaged using the GFP filter set, and PI was imaged using the RFP filter set. Three independent repeats were conducted.

### Starch staining assay

Flowers at each stage were collected at 4 hours after lights on. Samples were fixed in Carnoy fixative (ethanol:acetic acid, 3:1, v/v) overnight before moving to 70% ethanol and stored at 4°C. After sepals and petals were removed, extra ethanol was absorbed using tissue wipes. Samples were stained using 40 μl of clearing-staining solution [Image-iT™ Plant Tissue Clearing Reagent (Invitrogen, Waltham, MA, USA) with 4% (w/v) potassium iodide and 1.27% (w/v) iodine] before placing the cover slip. Samples were imaged by Zeiss Apotome.2 (Carl Zeiss, Thornwood, NY, USA) microscopy. More than 6 flowers at each stage were assayed for one repeat, and four independent repeats were conducted. For starch staining on wax embedded cross-sections, after stage 12 flower buds were fixed in Carnoy fixative overnight before moving to 70% ethanol, samples were dehydrated and embedded in wax as previously described (Zhang *et al*., 2021). Cross-sections (after wax was removed by HistoChoice^®^ Clearing Agent) were stained using 100 μl potassium iodide solution [composed of 4% (w/v) potassium iodide and 1.27% (w/v) iodine] kept in the dark for 5 min and washed twice using ddH2O before being imaged with a compound microscope (Nikon, NY, United States).

### Statistical analysis

The differences between the two subjects were determined using the two-tailed Student’s t-test with equal variance. The differences among multiple subjects were assessed using one-way ANOVA followed by multiple comparison tests (Fisher’s LSD method). All statistical analysis was performed using Origin 2021b statistical software (OriginLab Corporation, MA, USA).

### Accession numbers

Sequence information from this article can be found in the Arabidopsis Genome Initiative or GenBank/EMBL databases under the following accession numbers: AtSWEET9 (AT2G39060), AtSWEET13 (AT5G50800), AtSWEET14 (AT4G25010).

## Results

### AtSWEET13 and AtSWEET14 expressed at late anther development stages

*AtSWEET13* and *AtSWEET14* transcripts were detected in stamens at late-stage flower development stages using β-glucuronidase (GUS) reporter under the control of either *AtSWEET13* or *AtSWEET14* promoter (Kanno *et al*., 2016), but which exact anther development stages were yet to be determined. We examined the spatial and temporal protein accumulation of AtSWEET13 and AtSWEET14 using transgenic lines harboring a AtSWEET13 or AtSWEET14 translational fusion with GUS or with a yellow fluorescent protein (YFP) driven by its own native AtSWEET13 or AtSWEET14 promoter (named as *pSWEET13:SWEET13-GUS, pSWEET14:SWEET14-GUS, pSWEET13:SWEET13-YFP*, and *pSWEET14:SWEET14-YFP*). Both SWEET13-GUS protein and SWEET14-GUS were detected at late anther development stages in both unopened flower buds (anther development stage 12 and below) and fully opened flowers (anther development stages 13) (**Fig. 1a**) (Sanders *et al*., 1999). AtSWEET14 was also detected at the silique and pedicel junctions after fertilization, suggesting it may be involved in sugar transport from the pedicel to the silique. To determine at which anther development stage AtSWEET13 and AtSWEET14 start to function, we conducted the GUS staining of inflorescence tissues from *pSWEET13:SWEET13-GUS* and *pSWEET14:SWEET14-GUS* and embedded them in wax for cross-sectioning examination. Both SWEET13-GUS and SWEET14-GUS proteins were not detected at stage 11 (**Fig. 1b**), at which the tapetum is not fully degenerated, and pollen grains can be easily observed (Sanders *et al*., 1999). At stage 13 when the flower fully opens and pollen dehiscence occurs, both AtSWEET13 and AtSWEET14 were strongly accumulated in connective, epidermis, and endothecium cells (**Fig. 1b**). All stages stated here were confirmed by examining the flowers using *pSWEET13:SWEET13-YFP* and *pSWEET14:SWEET14-YFP* transgenic lines based on defined anther development stages (Bowman, 1994). Both AtSWEET13 and AtSWEET14 were only strongly detected at flower stages 12 and 13, corresponding to anther stages 11 to 13 (Sanders *et al*., 1999), and there were no signals observed in released pollen grains at anther stage 13 for both AtSWEET13 and AtSWEET14 (**Fig. 1c**). The anthers at stage 12 were further examined under confocal microscopy, AtSWEET13 and AtSWEET14 were strongly detected in the anther wall, including endothecium and epidermis cells (**Fig. 1d**), consistent with the detected GUS signal. In short, AtSWEET13 and AtSWEET14 were mainly expressed in connective, epidermis, and endothecium cells at anther development stages 12 and 13.

**Fig. 1.**
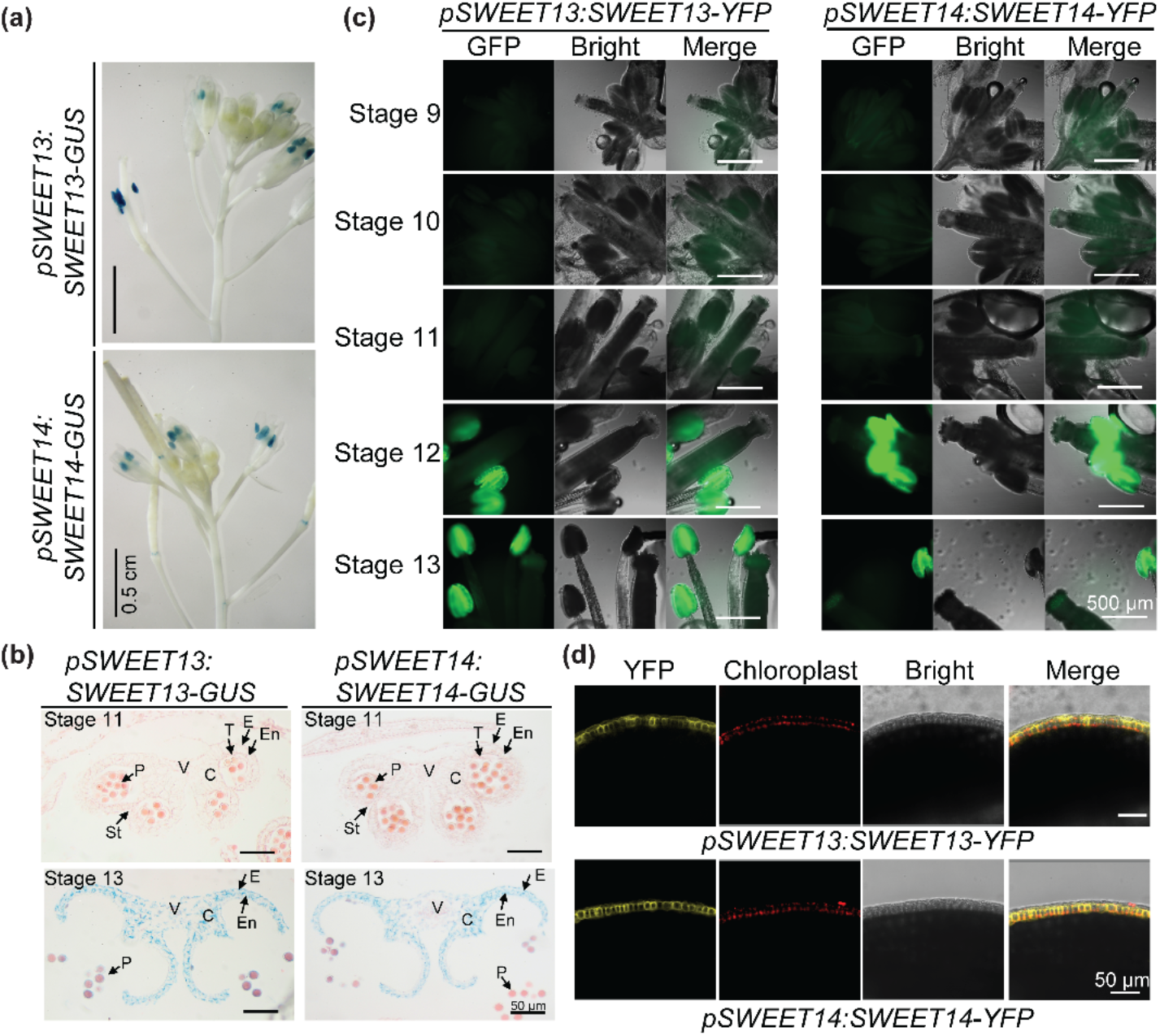
Spatial and temporal localizations of AtSWEET13 and AtSWEET14. **(a)** Tissue-specific expression of AtSWEET13 and AtSWEET14 was evaluated. Inflorescences from five-week-old Arabidopsis plants carrying *pSWEET13:SWEET13-GUS* and *pSWEET14:SWEET14-GUS* were histochemically stained for 12 h for GUS activity analysis. GUS staining patterns were consistent from two independent T2 lines for each construct. **(b)** Cross-sections of GUS-stained anther samples from *pSWEET13:SWEET13-GUS* and *pSWEET14:SWEET14-GUS* lines. C, connective; E, epidermis; En, endothecium; P, pollen grains; St, stomium; T, tapetum; V, vascular region. Section thickness: 6 μm. **(c)** AtSWEET13 and AtSWEET14 accumulation in different floral stages of Arabidopsis was examined using a fluorescence microscope. The signals were detected in anthers at floral stages 12 and 13 of transgenic lines carrying *pSWEET13:SWEET13-YFP* and *pSWEET14:SWEET14-YFP*. **(d)** AtSWEET13 and AtSWEET14 accumulation in anther wall of transgenic lines carrying *pSWEET13:SWEET13-YFP* and *pSWEET14:SWEET14-YFP* under confocal microscope. Stage 12 anthers were examined. Two independent lines were examined, and representative pictures from one line were shown.

### The reduced fertility phenotype of *atsweet13;14* is due to defected pollen

The double mutant of *atsweet13;14* has been reported to show a reduced fertility phenotype (Kanno *et al*., 2016). First, we confirmed the T-DNA insertion sites of *atsweet13* and *atsweet14* each single mutant was located in the second to last exon, and no full length *AtSWEET13* and *AtSWEET14* transcripts were detected in stage 13 flowers of *atsweet13;14* using RT-PCR (Supporting Information **Fig. S1**). As AtSWEET13 and AtSWEET14 were highly accumulated in anther, we speculated that the paternal effect on this phenotype is dominant. To test this, we reciprocally crossed *atsweet13;14* with Col-0. As shown in **Fig. 2(a)**, reduced fertility was exhibited when *atsweet13;14* pollen was crossed with Col-0 stigma or *atsweet13;14* stigma, while a normal seed set was observed when *atsweet13;14* stigma was crossed with Col-0 pollen. These data suggest that the reduced fertility phenotype of *atsweet13;14* is due to defected pollen. We further investigated whether the affected pollen is due to a defect in pollen viability or pollen germination. Pollen viability of *atsweet13;14*, was severely reduced compared to Col-0 (**Fig. 2b**), as demonstrated by fluorescein diacetate (FDA) and propidium iodide (PI) staining since viable pollen can be stained green by FDA and nonviable pollen can be stained red by PI. The defective *atsweet13;14* pollen viability can be fully complemented by either AtSWEET13 or AtSWEET14 under their own native promoters, as represented from two independent homozygous complementation lines (**Fig. 2b,c**). Around 65% of Col-0 pollen was alive, but less than 3% of *atsweet13;14* pollen was alive (**Fig. 2c**). Subsequently, *in vitro* pollen germination of the *atsweet13;14* pollen was analyzed. Similar to the pollen viability assay, pollen germination rate of *atsweet13;14* was severely reduced compared to Col-0 (**Fig. 2d**). Around 60% of Col-0 pollen germinated *in vitro*, while only 1% of *atsweet13;14* germinated (**Fig. 2e**). The defected *atsweet13;14* pollen germination can also be fully complemented by AtSWEET13 or AtSWEET14 under their own native promoters, as represented from two independent homozygous complementation lines (**Fig. 2d,e**). Thus, the reduced fertility phenotype of *atsweet13;14* is due to low pollen viability and in turn low pollen germination.

**Fig. 2.**
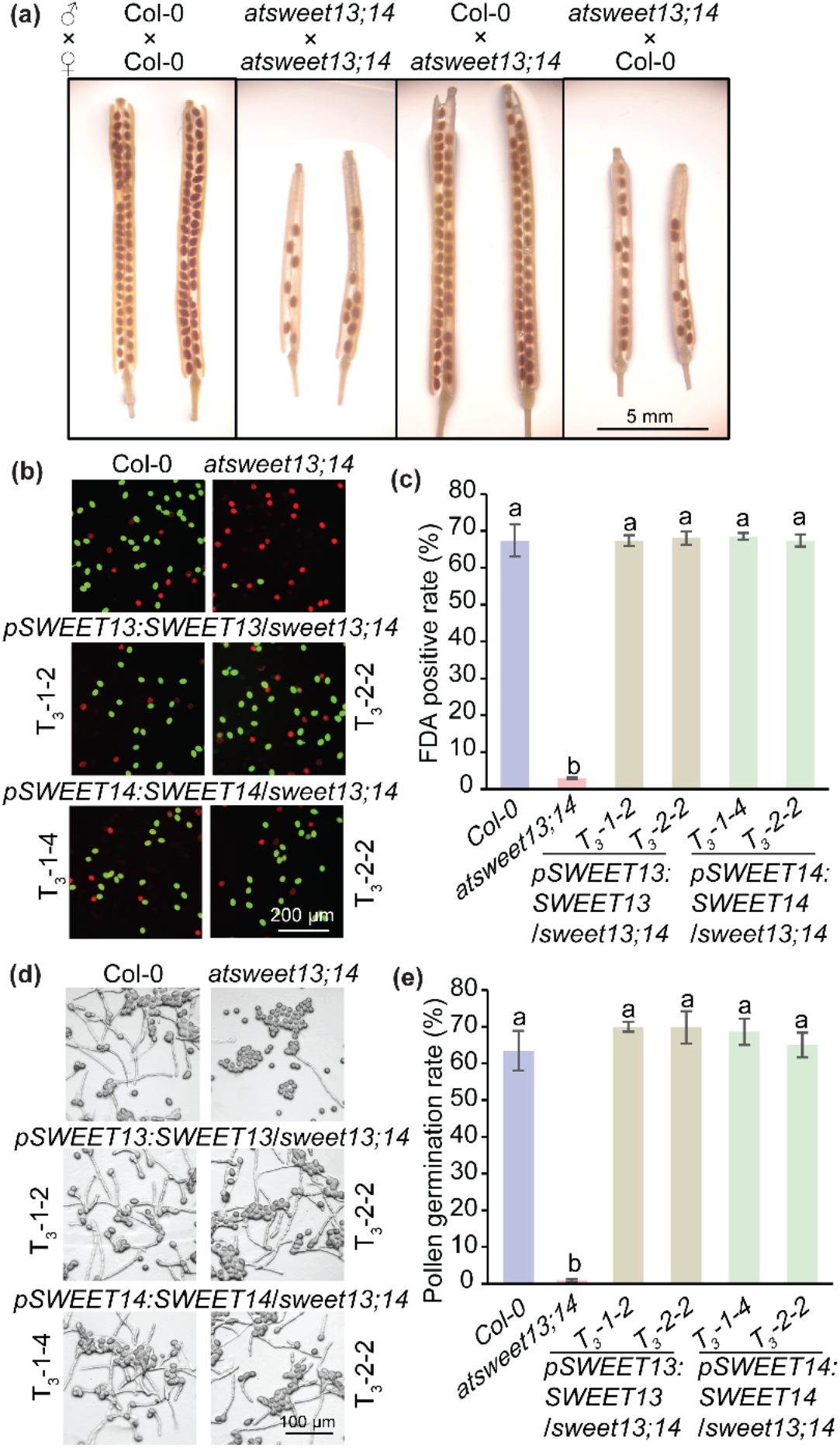
*atsweet13;14* pollen showed low viability and barely germinated *in vitro*. **(a)** Reciprocal crossing using pollen of Col-0 or *atsweet13;14* with stigma of Col-0 or *atsweet13;14*. 14-day-old siliques were imaged using a dissecting microscope. **(b)** FDA and PI staining of pollen from Col-0, *atsweet13;14*, and complementation lines. Pollen was collected directly from fresh flowers. Live pollen was stained green by FDA, and dead pollen was stained red by PI. **(c)** Statistical analysis of FDA positive rates of pollen from Col-0 and *atsweet13;14*, and complementation lines. The means were calculated from multiple repeats (± SE, n = 6), with over 900 pollen grains counted in total. **(d)** *In vitro* pollen germination analysis of Col-0, *atsweet13;14* and complementation lines. The pictures were taken 8 hours post germination. **(e)** Statistical analysis of pollen germination rates for various genotypes. The means were calculated from multiple repeats (± SE, n = 6), with over 450 pollen grains/tubes counted in total. The statistically significant differences among different samples in panel (c) and (e) were determined using one-way ANOVA followed by multiple comparison tests (LSD method) and were represented by different letters (*P* <0.05).

### Disrupted sucrose transport is responsible for the defective *atsweet13;14* pollen fertility

Both sucrose and GA have been demonstrated as the substrates of AtSWEET13 or AtSWEET14 (Chen *et al*., 2012; Kanno *et al*., 2016). To address which substrate is responsible for the defected *atsweet13;14* pollen, we used *AtSWEET14* promoter to drive AtSWEET9 to complement *atsweet13;14*, since AtSWEET9 transports sucrose but not GA. We first confirmed SWEET9-YFP was detected in the anthers of flowers at stages 12 and 13 (Supporting Information **Fig. S2**), and further confirmed SWEET9-YFP was accumulated in the anther wall at anther stage 12 (**Fig. 3a**), similar to the temporospatial accumulation of either SWEET13-YFP or SWEET14-YFP under its own promoter (**Fig. 1c,d**). As a result, AtSWEET9 fully complemented *atsweet13;14* pollen viability (**Fig. 3b,c**) and germination phenotype (**Fig. 3d,e**), as represented from two independent homozygous complementation lines. These data support that sucrose transport mediated by AtSWEET13 and AtSWEET14 is essential for supporting pollen viability and subsequent pollen germination.

**Fig. 3.**
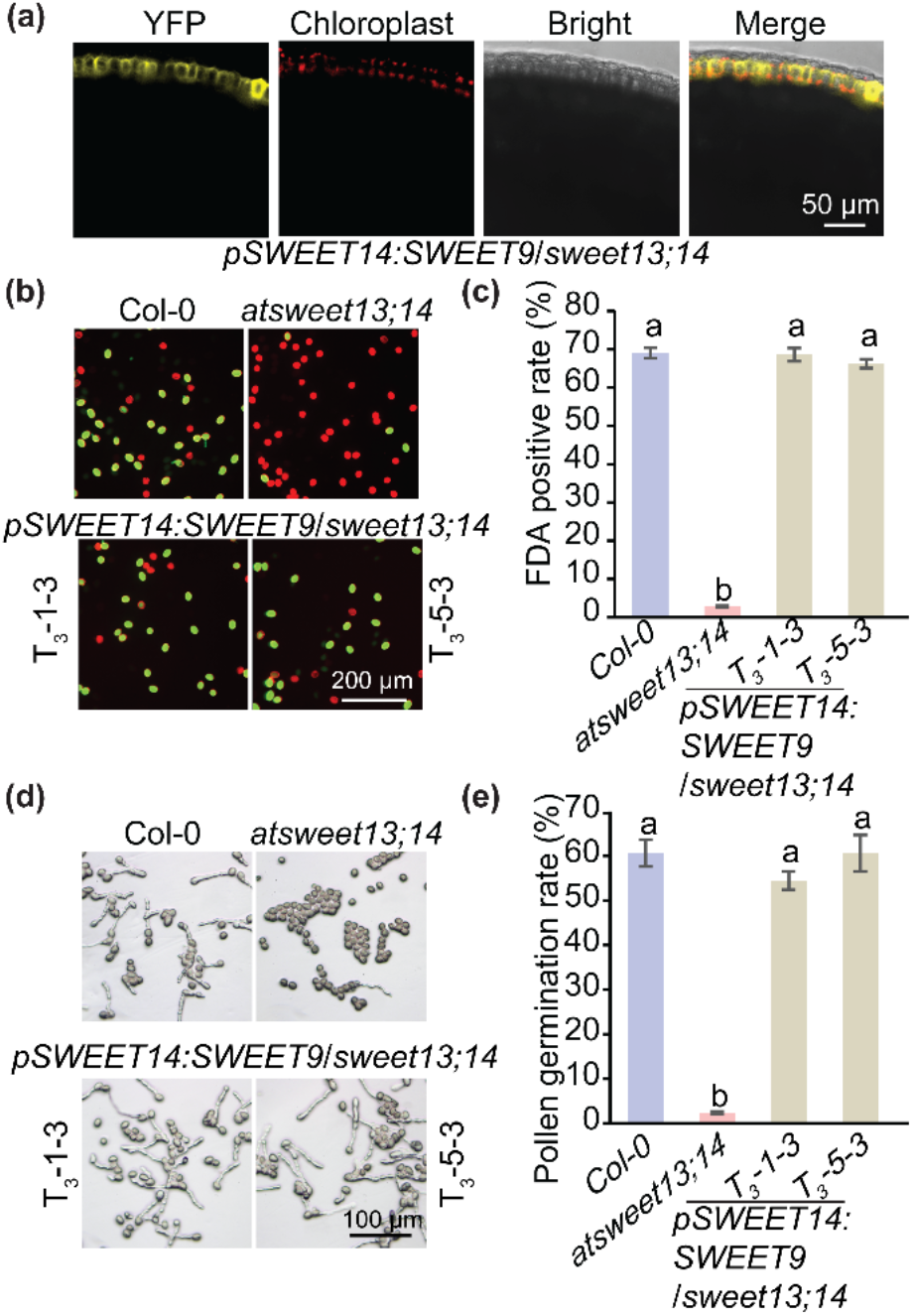
Defected pollen of *atsweet13;14* was rescued by AtSWEET9. **(a)** AtSWEET9 accumulation in anther wall of *atsweet13;14* complementation lines carrying *pSWEET14:SWEET9-YFP* under confocal microscope. Stage 12 anthers of *atsweet13;14* complementation lines were examined. Two independent complementation lines were examined, and representative pictures from one line were shown. **(b)** Complementation analysis of pollen viability of *atsweet13;14* by *pSWEET14:SWEET9-YFP*. AtSWEET9 rescued low pollen viability of *atsweet13;14*. Live pollen was stained green by FDA, and dead pollen was stained red by PI. **(c)** Statistical analysis of FDA positive rates for AtSWEET9 complementation lines. The means were calculated from multiple repeats (± SE, n = 12), with over 1400 pollen grains counted in total. **(d)** Complementation analysis of *in vitro* pollen germination of *atsweet13;14* by *pSWEET14:SWEET9-YFP*. AtSWEET9 rescued low pollen germination of *atsweet13;14*. The pictures were taken 8 hours post germination. **(e)** Statistical analysis of pollen germination rates for AtSWEET9 complementation lines. The means were calculated from multiple repeats (± SE, n = 6), with over 300 pollen grains/tubes counted in total. The statistically significant differences among different samples in panel **(d)** and **(f)** were determined using one-way ANOVA followed by multiple comparison tests (LSD method) and were represented by different letters (*P* <0.05).

### AtSWEET13 and AtSWEET14 function as gatekeepers to regulate sugar distribution between anther and pollen at late anther stages

Pollen is symplasmically isolated from the somatic cell layers of anther wall where AtSWEET13 and AtSWEET14 proteins were located. Sucrose is likely to be exported by AtSWEET13 and AtSWEET14 from the endothecium into locules before being taken up by pollen. We speculated that more sugar would accumulate in the anther wall of *atsweet13;14* if the endothecium was the site to export sugar. We have also learned that sugar availability is tightly associated with starch accumulation, and starch dynamics have been observed in the stamen envelope (Hedhly *et al*., 2016). Thus, instead of measuring soluble sugar content directly from stamen cell walls, which are more challenging, we compared starch staining between Col-0 and *atsweet13;14*. As expected, more starch accumulation was observed in *atsweet13;14* anthers at stage 12 than that in Col-0 (Supporting Information **Fig. S3**). To confirm the differential starch accumulation at anther stage 12, we embedded flower buds of Col-0 and *atsweet13;14* in wax for cross-sectioning examination followed by starch staining. The substantial starch granules were clearly observed in connective, epidermis, and endothecium cells of *atsweet13;14* compared with Col-0 at anther stage 12 (**Fig. 4a,b**). Although there was higher starch with an indication of high soluble sugar in the anther wall of *atsweet13;14*, no structural differences of anthers were observed when scanned under a scanning electron microscope (SEM) at stage 12 and stage 13 (**Fig. 4c,d**). Consistent with the reported observation (Kanno *et al*., 2016), *atsweet13;14* showed delayed anther dehiscence at stage 12 (**Fig. 4c**). Unsurprisingly, we found the size of *atsweet13;14* pollen was affected as demonstrated by the significantly reduced length and width compared to Col-0 pollen at both stages 12 and 13 (**Fig. 4e,f**), likely resulting from the lack of sugar supply from the anther wall, which is consistent with the low pollen viability (**Fig. 2b,c**). Notably, the width of Col-0 pollen was significantly decreased from stage 12 to stage 13 (**Fig. 4f**), in agreement with the reported dehydration process taken by Arabidopsis pollen grain at anthesis (Shi & Yang, 2010). To confirm that insufficient sugar supply is likely the cause of pollen defects, we measured soluble sugar levels from the mature pollen. All sugars, including sucrose (**Fig. 4g**), glucose (**Fig. 4h**), and fructose (**Fig. 4i**), substantially decreased in *atsweet13;14* pollen compared to Col-0 pollen. Lipid is another major form of carbon storage in mature pollen grains, especially for Arabidopsis (Shi & Yang, 2010; Wang *et al*., 2022a). Sugar is the primary source for lipid body biogenesis in pollen grains. Defects in lipid body accumulation affect pollen fitness and successful fertilization (Zheng *et al*., 2018). Thus, we wondered whether the total fatty acids altered, which in turn could contribute to the low pollen germination in *atsweet13;14*. Surprisingly, the total fatty acids in *atsweet13;14* remained similar to that in Col-0 (**Fig. 4j**). In addition, no significant differences were observed in each fatty acids species commonly found in Arabidopsis pollen (Wang *et al*., 2022a), including C16:0, C18:0, C18:1, C18:2, and C18:3 (Supporting Information **Fig. S4**).

**Fig. 4.**
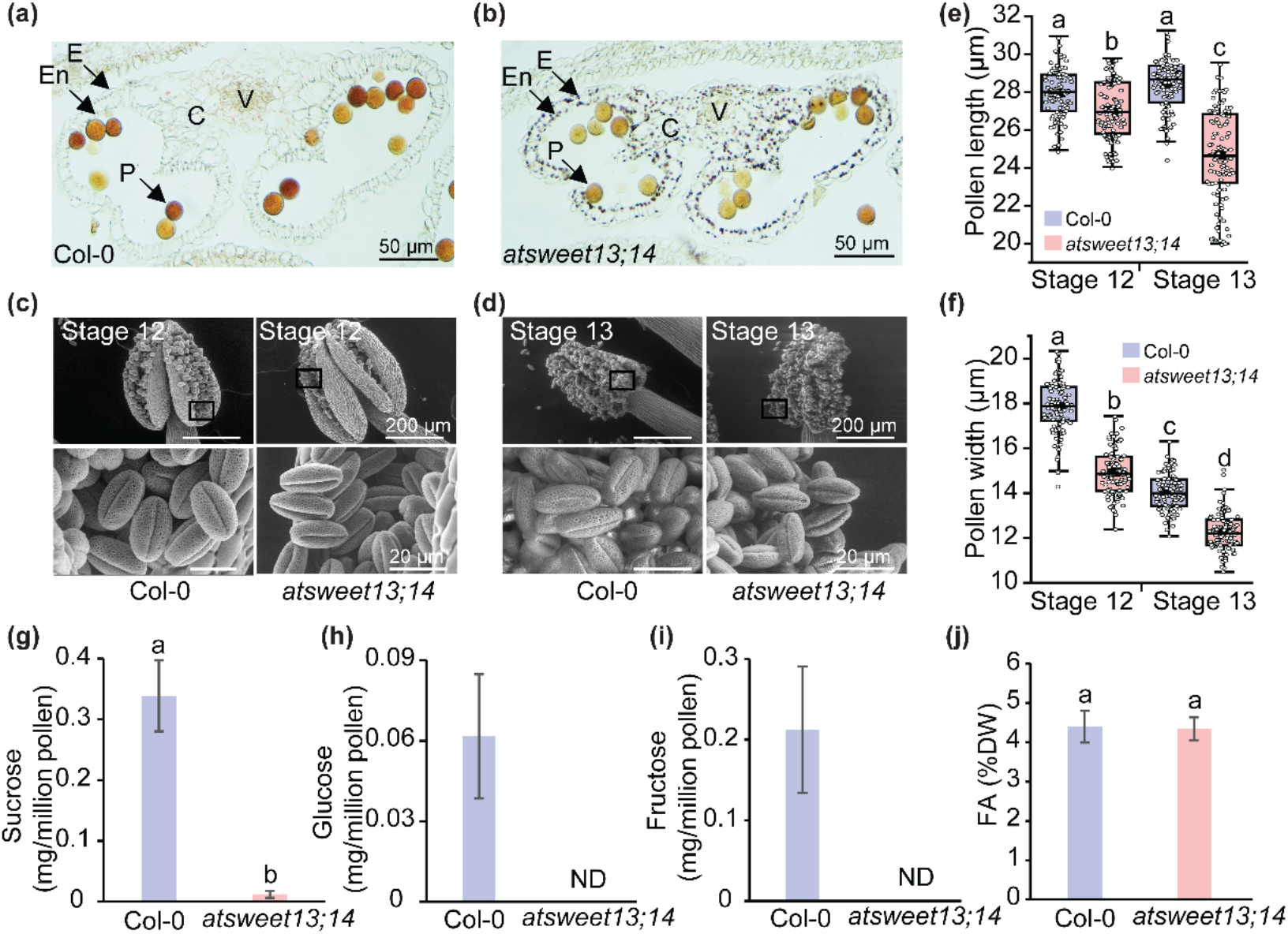
Phenotypic effects of *atsweet13;14* on anthers and pollen. **(a)** and **(b)** Comparison of starch accumulation in anther cross-sections between Col-0 and *atsweet13;14* at stage 12. Starch granules were accumulated in endothecium, epidermis, and connective cells of *atsweet13;14* anther compared with those in Col-0. C, connective; E, epidermis; En, endothecium; P, pollen grains; V, vascular region. Section thickness: 6 μm. **(c)** and **(d)** Comparison of anther and pollen morphological phenotypes imaged by SEM between Col-0 and *atsweet13;14* at stage 12 **(c)** and stage 13 **(d)**. The close-up pictures at the bottom were taken in the marked rectangle region from the top corresponding image. (e) and **(f)** Quantitative comparison of pollen size between Col-0 and *atsweet13;14* at stages 12 and 13. **(e)** Comparison in length and **(f)** comparison in width measured by Image J from 99 pollen grains with near flat orientation. The boxes were drawn from the 25^th^ percentile to the 75^th^ percentile; whiskers extended to 1.5× the interquartile range from the 25^th^ and 75^th^ percentiles; the medians were shown in line and the means were shown in black square. The statistical analysis was done using one-way ANOVA followed by LSD method and differences were represented by different letters (*P* <0.05). **(g-i)** Comparison of soluble sugar contents measured from Col-0 and *atsweet13;14* mutant pollen grains directly collected from fresh flowers. Sucrose **(g)**, glucose **(h)** and fructose **(i)** in pollen were quantified from fresh open flowers collected in three independent repeats (means ± SE, n = 3). Glucose and fructose content in *atsweet13;14* was below the detection limit using our established method. ND, not detected. **(j)** Comparison of total fatty acids content from Col-0 and *atsweet13;14* mutant pollen. Total fatty acids content of Col-0 and *atsweet13;14* pollen collected directly from fresh flowers (means ± SE, n = 3). Fatty acid methyl esters (FAME) extracted from four independent repeats (± SE, n = 4) were quantified using GC-FID. The statistically significant differences in panel (**g** and **j**) were determined using Student’s t-test and were represented by different letters (*P* <0.05).

## Discussion

### Pollen fertility phenotypes are explained by sucrose rather than GA as a substrate of AtSWEET13 and AtSWEET14

Transport proteins are essential for cells to selectively exchange solutes across the cell membrane in many physiological processes, including nutrient transport. It is not uncommon for a single transporter to transport multiple substrates that are structurally distant. For example, AtGTR1/NPF2.10 transports glucosinolates, JA-Ile and GA (Saito *et al*., 2015); AtNPF3.1/NRT1.1 transports nitrate, GA, and abscisic acid (ABA) (Tal *et al*., 2016; David *et al*., 2016); AtNPF6.3 transports both nitrate and auxin (Krouk *et al*., 2010); AtNPF4.1/AIT3 transports both ABA and GA (Kanno *et al*., 2012); AtPHT4.4 transports phosphate and ascorbate (Guo *et al*., 2008; Miyaji *et al*., 2015); AtABCG36 transports indole-3-butyric acid and cadmium (Strader & Bartel, 2009); A classical sucrose/H^+^ symporter AtSUC5 transports biotin in addition to sucrose (Ludwig *et al*., 2000), and this was the first report of a sugar transporter to transport a structurally unrelated substrate from sugar.

It has been over a decade since SWEETs have been characterized as bidirectional uniporters transporting sugars (Chen *et al*., 2010; Xue *et al*., 2022). In 2016, Kanno and co-authors reported that AtSWEET10, AtSWEET12, AtSWEET13 and AtSWEET14 were able to mediate GA transport using a modified yeast two-hybrid system containing BD-GID1a (GA receptor) and AD-GAI (a DELLA protein). The same system has been used to detect weak GA transport activity for OsSWEET3a, OsSWEET11a and OsSWEET12 (Kanno *et al*., 2016; Morii *et al*., 2020; Wu *et al*., 2022). However, the biological relevance of GA transport mediated by SWEETs was only indicated by partial restoration of anther dehiscence of the *atsweet13;14* mutant through exogenous GA3 application (Kanno *et al*., 2016), and restoration of defects in germination and early shoot development for *ossweet3a* (Morii *et al*., 2020).

In addition, only a few members of SWEETs from Arabidopsis and rice transport GA, offering a chance to study the preferred substrate for a given biological process. Kanno et al. (2016) indicated that AtSWEET13 and AtSWEET14 mediated GA transport in the anther. AtSWEET9 and AtSWEET12 show a similar sucrose transport activity to AtSWEET13 and AtSWEET14 in a biosensor-based HEK293T detection system (Chen *et al*., 2012), but AtSWEET9 does not show any GA transport activity (Kanno *et al*., 2016). In addition, AtSWEET12, under the control of the AtSWEET9 promoter, is sufficient to rescue the defect in the nectar secretion phenotype of the *atsweet9* mutant, further supporting the conclusion that they are exchangeable in a physiological context when expressed at the same site (Lin *et al*., 2014). Following this concept, we generated transgenic plants with *AtSWEET9* expression driven by the promoter of *AtSWEET14* and found this combination can fully complement the compromised pollen germination phenotype of *atsweet13;14* (**Figure 3**). Moreover, the soluble sugar content (including sucrose, glucose and fructose) was dramatically reduced in pollen of *atsweet13;14* compared with Col-0 (**Fig.4g,h,i**), while the content of all measured GA species was not significantly affected in anthers between Col-0 and *atsweet13;14* (Kanno *et al*., 2016). Notably, one parallel study shows that the sucrose selective AtSWEET13 mutant was able to fully rescue the pollen viability and germination phenotype of *atsweet13;14* but GA selective AtSWEET13 mutant was not (Isoda submitted), which is consistent with our findings.

### Endothecium is the major site where sucrose is transported by AtSWEET13 and AtSWEET14

Pollen development requires nutrient supply from locules of the anthers. The question of how sugars reach locules is poorly understood. The anther structure changes dynamically in a development-dependent manner. The anther wall has four layers, i.e., the epidermis, endothecium, middle layers and tapetum. The epidermis plays a protective role during the anther development. The endothecium, likely serving as an energy storage tissue, stores starch and is associated with anther dehiscence for pollen release (van der Linde & Walbot, 2019). Little is known about the function of the middle layer, which disappears entirely at stage 11 (Xue *et al*., 2021). The tapetum facilitates the flow of nutrients and water for microspore development and secretes exine components for pollen formation (van der Linde & Walbot, 2019). The tapetum initiates degeneration at stage 10 and completely disappears at stage 12 (Sanders *et al*., 1999). Our results showed that AtSWEET13 and AtSWEET14 proteins were detected at stage 12 and stage 13 (**Fig. 1b,c**) when both the middle layer and tapetum degenerated. AtSWEEET13-GUS/YFP and AtSWEET14-GUS/YFP were mainly observed in the wall of the anther (**Fig.1b,d**). These data suggest sucrose efflux mediated by AtSWEET13 and AtSWEET14 from endothecium cells into the cavity of the anther mainly occurs at stages 12 and 13, when the endothecium is the innermost layer of the anther wall. Thus, it was reasonable to predict that impaired sucrose efflux from the endothecium in *atsweet13;14* mutant can lead to higher sucrose accumulation and in turn more starch would be synthesized in the anther, which was demonstrated by the excess starch accumulation in connective, epidermis, and endothecium cells of *atsweet13;14* at stage 12 (**Fig. 4a,b**). Consistently, pollen grains were detected to have a lower level of sugar level due to less sugar being available from the wall of the anther in *atsweet13;14*. Although pollen viability and pollen germination were dramatically reduced, there still was a low rate of alive pollen in *atsweet13;14*, which may be due to the expression of clade III *AtSWEETs* (Chen *et al*., 2015; Mergner *et al*., 2020) or the degeneration of starch synthesized in pollen from earlier developmental stages (Hedhly *et al*., 2016).

Interestingly, the total fatty acid content of *atsweet13;14* remains comparable with that of Col-0 (**Fig. 4j**, Supporting Information **Fig. S4**), which could be synthesized before the stages when AtSWEET13 and AtSWEET14 start to function during the anther development. Tapetum accumulates lipidic components for pollen coat formation (Piffanelli *et al*., 1998; Ariizumi *et al*., 2004), while *AtSWEET13* and *AtSWEET14* are highly expressed after tapetum degeneration, which indicates mature pollen grains have received some lipidic components from tapetum even in *atsweet13;14*. In addition, six sugar transporters are expressed at earlier stages of anther development (Feng *et al*., 2012), which may provide pollen with sugar for carbon storage. mRNA encoding a functional acyl-CoA:diacylglycerol acyltransferase (AtDGAT2) (Zhou *et al*., 2013) is strongly transcribed in Arabidopsis microspore before gradually decreasing in mature pollen grains (Honys & Twell, 2004). The carbon needed for fatty acid synthesis at early anther stages is likely through tapetum-expressed sugar transporters, such as *AtSWEET8* (Guan *et al*., 2008). Sucrose transporter AtSWEET15 is also accumulated in the anthers of early buds (Chen *et al*., 2015). This possible explanation is also supported by the observation that lipid bodies were already visible in microspores of tobacco pollen (Rotsch *et al*., 2017). Similarly, the total fatty acid content of *atsuc1* pollen was also comparable to that of Col-0, though the pollen germination of *atsuc1* was substantially compromised (Sivitz *et al*., 2008). Further investigation is needed to elucidate the underlying mechanism. It would be most helpful to conduct metabolite analysis of soluble sugar, amino-acid, and fatty acid contents across pollen development stages is performed in Arabidopsis, which produces tricellular pollen, different from the bicellular pollen from tobacco. In tobacco, sucrose content increases substantially right before pollen matures and peaks in mature pollen grains (Rotsch *et al*., 2017), which could be due to increased sucrose unloading and/or reduced storage conversion. It is worth addressing whether any SWEETs, such as homologs of AtSWEET13 and AtSWEET14 are involved in the increased sucrose levels.

To summarize, we proposed a model to illustrate the role of sugar transporters during the late anther developmental stage (**Fig. 5**). Photoassimilates are symplasmically unloaded from the phloem of the filament to the anther connective tissue (Imlau *et al*., 1999). AtSWEET13 and AtSWEET14 proteins are found in the connective tissue, epidermis, and endothecium at late anther stage (**Fig. 1b**) and participate in sucrose apoplasmic unloading into the locules. The released sucrose exported by AtSWEET13 and AtSWEET14 is likely to be partially cleaved into glucose and fructose by cell-wall invertase 2 (AtcwINV2) or other AtcwINVs, because the *AtcwINV2 antisense* mutant displays a defective pollen and reduced fertility phenotype similar to *atsweet13;14* (Hirsche *et al*., 2009). The resulting hexoses and sucrose may be taken up by pollen grain-accumulated AtSWEET5 (Ko *et al*., 2022; Wang *et al*., 2022b), AtSWEET8 (Zhang *et al*., 2022) and AtSUC1 (Stadler *et al*., 1999; Sivitz *et al*., 2008), respectively.

**Fig. 5.**
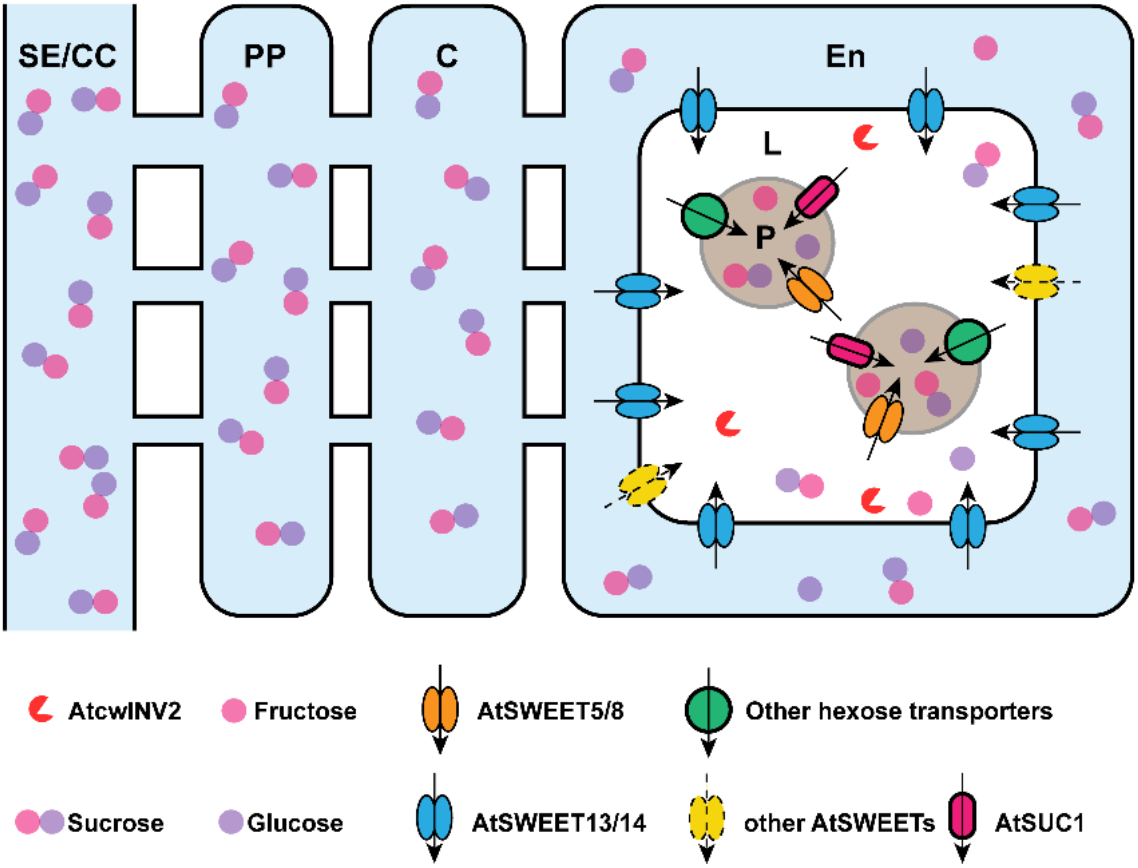
Schematic representation of the role of AtSWEET13/14 during apoplasmic unloading at anther stage12. SE, sieve element; CC, companion cells; C, connective cells; En, endothecium; P, pollen grains; L, locule.

Overall, we concluded that the impaired pollen viability is responsible for the reduced fertility of *atsweet13;14*. Sucrose transported by AtSWEET13 and AtSWEET14 is responsible for the defective pollen phenotype of *atsweet13;14*. AtSWEET13 and AtSWEET14 mainly export sucrose from anther wall to locules.

## Acknowledgement

We thank Dr. Yaxin Li for the help on soluble sugar measurement, we also thank Catherine Wallace for the help on SEM imaging. This work is supported by startup funds from the University of Illinois at Urbana-Champaign to Dr. Li-Qing Chen (to JW, XX, JL). HZ was supported by a scholarship from the China Scholarship Council (201708330168).

## Author contributions

L-QC oversaw the whole project. JW, XX, and L-QC designed the work. JW, XX, HZ and JL conducted the experiments. JW, XX and HZ analyzed the data. JW and L-QC wrote the manuscript.

## Data availability

All the data and materials that support the findings of this study are available upon request from the corresponding author.

## Supporting Information

Additional supporting information may be found in the online version of this article.

**Fig. S1** Genotyping of *atsweet13;14* double mutant.

**Fig. S2** AtSWEET9 accumulation in different floral stages of Arabidopsis was examined using fluorescence microscopy.

**Fig. S3** Comparison of starch accumulation in anthers between Col-0 and *atsweet13;14* at various stages.

**Fig. S4** Comparison of fatty acid species between Col-0 and *atsweet13;14* pollen.

**Table S1** Primers used in this study.

## References

Ariizumi T, Hatakeyama K, Hinata K, Inatsugi R, Nishida I, Sato S, Kato T, Tabata S, Toriyama K. 2004. Disruption of the novel plant protein NEF1 affects lipid accumulation in the plastids of the tapetum and exine formation of pollen, resulting in male sterility in Arabidopsis thaliana. The Plant Journal 39: 170–181.

Binenbaum J, Weinstain R, Shani E. 2018. Gibberellin localization and transport in plants. Trends in Plant Science 23: 410–421.

Borghi M, Fernie AR. 2017. Floral metabolism of sugars and amino acids: implications for pollinators’ preferences and seed and fruit set. Plant Physiology 175: 1510–1524.

Bowman J. 1994. Arabidopsis: an atlas of morphology and development.

Braun DM. 2022. Phloem loading and unloading of sucrose: what a long, strange trip from source to sink. Annual Review of Plant Biology 73: in press.

Browse J, McCourt PJ, Somerville CR. 1986. Fatty acid composition of leaf lipids determined after combined digestion and fatty acid methyl ester formation from fresh tissue. Analytical Biochemistry 152: 141–145.

Chen L-Q, Hou B-H, Lalonde S, Takanaga H, Hartung ML, Qu X-Q, Guo W-J, Kim J-G, Underwood W, Chaudhuri B, et al. 2010. Sugar transporters for intercellular exchange and nutrition of pathogens. Nature 468: 527–532.

Chen L-Q, Lin IW, Qu X-Q, Sosso D, McFarlane HE, Londoño A, Samuels AL, Frommer WB. 2015. A cascade of sequentially expressed sucrose transporters in the seed coat and endosperm provides nutrition for the Arabidopsis embryo. The Plant Cell 27: 607–619.

Chen L-Q, Qu X-Q, Hou B-H, Sosso D, Osorio S, Fernie AR, Frommer WB. 2012. Sucrose efflux mediated by SWEET proteins as a key step for phloem transport. Science 335: 207–211.

Clément C, Audran JC. 1995. Anther wall layers control pollen sugar nutrition in Lilium. Protoplasma 187: 172–181.

Clough SJ, Bent AF. 1998. Floral dip: a simplified method for Agrobacterium-mediated transformation of *Arabidopsis thaliana*. The Plant Journal 16: 735–743.

David LC, Berquin P, Kanno Y, Seo M, Daniel-Vedele F, Ferrario-Méry S. 2016. N availability modulates the role of NPF3.1, a gibberellin transporter, in GA-mediated phenotypes in Arabidopsis. Planta 244: 1315–1328.

Feng B, Lu D, Ma X, Peng Y, Sun Y, Ning G, Ma H. 2012. Regulation of the Arabidopsis anther transcriptome by DYT1 for pollen development. The Plant Journal 72: 612–624.

Griffiths J, Murase K, Rieu I, Zentella R, Zhang Z-L, Powers SJ, Gong F, Phillips AL, Hedden P, Sun T, et al. 2006. Genetic characterization and functional analysis of the GID1 gibberellin receptors in Arabidopsis. The Plant Cell 18: 3399–3414.

Guan Y-F, Huang X-Y, Zhu J, Gao J-F, Zhang H-X, Yang Z-N. 2008. RUPTURED POLLEN GRAIN1, a member of the MtN3/saliva gene family, is crucial for exine pattern formation and cell integrity of microspores in Arabidopsis. Plant Physiology 147: 852–863.

Guo B, Jin Y, Wussler C, Blancaflor EB, Motes CM, Versaw WK. 2008. Functional analysis of the Arabidopsis PHT4 family of intracellular phosphate transporters. New Phytologist 177: 889–898.

Hedhly A, Vogler H, Schmid MW, Pazmino D, Gagliardini V, Santelia D, Grossniklaus U. 2016. Starch turnover and metabolism during flower and early embryo development. Plant Physiology 172: 2388–2402.

Hirose T, Zhang Z, Miyao A, Hirochika H, Ohsugi R, Terao T. 2010. Disruption of a gene for rice sucrose transporter, OsSUT1, impairs pollen function but pollen maturation is unaffected. Journal of Experimental Botany 61: 3639–3646.

Hirsche J, Engelke T, Völler D, Götz M, Roitsch T. 2009. Interspecies compatibility of the anther specific cell wall invertase promoters from Arabidopsis and tobacco for generating male sterile plants. Theoretical and Applied Genetics 118: 235–245.

Honys D, Twell D. 2004. Transcriptome analysis of haploid male gametophyte development in Arabidopsis. Genome Biology 5: R85.

Imlau A, Truernit E, Sauer N. 1999. Cell-to-cell and long-distance trafficking of the green fluorescent protein in the phloem and symplastic unloading of the protein into sink tissues. The Plant Cell 11: 309–322.

Itoh H, Tanaka-Ueguchi M, Kawaide H, Chen X, Kamiya Y, Matsuoka M. 1999. The gene encoding tobacco gibberellin 3β-hydroxylase is expressed at the site of GA action during stem elongation and flower organ development. The Plant Journal 20: 15–24.

Kanno Y, Hanada A, Chiba Y, Ichikawa T, Nakazawa M, Matsui M, Koshiba T, Kamiya Y, Seo M. 2012. Identification of an abscisic acid transporter by functional screening using the receptor complex as a sensor. Proceedings of the National Academy of Sciences 109: 9653–9658.

Kanno Y, Oikawa T, Chiba Y, Ishimaru Y, Shimizu T, Sano N, Koshiba T, Kamiya Y, Ueda M, Seo M. 2016. AtSWEET13 and AtSWEET14 regulate gibberellin-mediated physiological processes. Nature Communications 7: 13245.

Ko H-Y, Tseng H-W, Ho L-H, Wang L, Chang T-F, Lin A, Ruan Y-L, Neuhaus HE, Guo W-J. 2022. Hexose translocation mediated by SlSWEET5b is required for pollen maturation in *Solanum lycopersicum*. Plant Physiology: kiac057.

Krouk G, Lacombe B, Bielach A, Perrine-Walker F, Malinska K, Mounier E, Hoyerova K, Tillard P, Leon S, Ljung K, et al. 2010. Nitrate-regulated auxin transport by NRT1.1 defines a mechanism for nutrient sensing in plants. Developmental Cell 18: 927–937.

Kubo M, Udagawa M, Nishikubo N, Horiguchi G, Yamaguchi M, Ito J, Mimura T, Fukuda H, Demura T. 2005. Transcription switches for protoxylem and metaxylem vessel formation. Genes & Development 19: 1855–1860.

Lin IW, Sosso D, Chen L-Q, Gase K, Kim S-G, Kessler D, Klinkenberg PM, Gorder MK, Hou B-H, Qu X-Q, et al. 2014. Nectar secretion requires sucrose phosphate synthases and the sugar transporter SWEET9. Nature 508: 546–549.

van der Linde K, Walbot V. 2019. Pre-meiotic anther development. In: Grossniklaus U, ed. Plant Development and Evolution. Current Topics in Developmental Biology. Academic Press, 239–256.

Ludwig A, Stolz J, Sauer N. 2000. Plant sucrose-H+ symporters mediate the transport of vitamin H. The Plant Journal 24: 503–509.

Ma H. 2005. Molecular genetic analyses of microsporogenesis and microgametogenesis in flowering plants. Annual Review of Plant Biology 56: 393–434.

Matsoukas IG. 2014. Interplay between sugar and hormone signaling pathways modulate floral signal transduction. Frontiers in Genetics 5.

Mergner J, Frejno M, List M, Papacek M, Chen X, Chaudhary A, Samaras P, Richter S, Shikata H, Messerer M, et al. 2020. Mass-spectrometry-based draft of the Arabidopsis proteome. Nature 579: 409–414.

Miyaji T, Kuromori T, Takeuchi Y, Yamaji N, Yokosho K, Shimazawa A, Sugimoto E, Omote H, Ma JF, Shinozaki K, et al. 2015. AtPHT4;4 is a chloroplast-localized ascorbate transporter in Arabidopsis. Nature Communications 6: 5928.

Morii M, Sugihara A, Takehara S, Kanno Y, Kawai K, Hobo T, Hattori M, Yoshimura H, Seo M, Ueguchi-Tanaka M. 2020. The dual function of OsSWEET3a as a gibberellin and glucose transporter is important for young shoot development in rice. Plant and Cell Physiology 61:1935–1945.

Muhlemann JK, Younts TLB, Muday GK. 2018. Flavonols control pollen tube growth and integrity by regulating ROS homeostasis during high-temperature stress. Proceedings of the National Academy of Sciences 115: E11188–E11197.

Piffanelli P, Ross JHE, Murphy DJ. 1998. Biogenesis and function of the lipidic structures of pollen grains. Sexual Plant Reproduction 11: 65–80.

Plackett ARG, Thomas SG, Wilson ZA, Hedden P. 2011. Gibberellin control of stamen development: a fertile field. Trends in Plant Science 16: 568–578.

Rotsch AH, Kopka J, Feussner I, Ischebeck T. 2017. Central metabolite and sterol profiling divides tobacco male gametophyte development and pollen tube growth into eight metabolic phases. The Plant Journal 92: 129–146.

Saito H, Oikawa T, Hamamoto S, Ishimaru Y, Kanamori-Sato M, Sasaki-Sekimoto Y, Utsumi T, Chen J, Kanno Y, Masuda S, et al. 2015. The jasmonate-responsive GTR1 transporter is required for gibberellin-mediated stamen development in Arabidopsis. Nature Communications 6: 6095.

Sanders PM, Bui AQ, Weterings K, McIntire KN, Hsu Y-C, Lee PY, Truong MT, Beals TP, Goldberg RB. 1999. Anther developmental defects in *Arabidopsis thaliana* male-sterile mutants. Sexual Plant Reproduction 11: 297–322.

Shi D, Yang W. 2010. Pollen germination and tube growth. Plant developmental biology-biotechnological perspectives: 245–282.

Sivitz AB, Reinders A, Ward JM. 2008. Arabidopsis sucrose transporter AtSUC1 is important for pollen germination and sucrose-induced anthocyanin accumulation. Plant Physiology 147: 92–100.

Stadler R, Truernit E, Gahrtz M, Sauer N. 1999. The AtSUC1 sucrose carrier may represent the osmotic driving force for anther dehiscence and pollen tube growth in Arabidopsis. Plant Journal 19: 269–278.

Strader LC, Bartel B. 2009. The Arabidopsis PLEIOTROPIC DRUG RESISTANCE8/ABCG36 ATP binding cassette transporter modulates sensitivity to the auxin precursor indole-3-butyric acid. The Plant Cell 21: 1992–2007.

Sun T, Goodman HM, Ausubel FM. 1992. Cloning the Arabidopsis GA1 locus by genomic subtraction. The Plant Cell 4: 119–128.

Sun M-X, Huang X-Y, Yang J, Guan Y-F, Yang Z-N. 2013. Arabidopsis RPG1 is important for primexine deposition and functions redundantly with RPG2 for plant fertility at the late reproductive stage. Plant Reproduction 26: 83–91.

Sun L, Sui X, Lucas WJ, Li Y, Feng S, Ma S, Fan J, Gao L, Zhang Z. 2019. Down-regulation of the sucrose transporter CsSUT1 causes male sterility by altering carbohydrate supply. Plant Physiology 180: 986–997.

Tal I, Zhang Y, Jørgensen ME, Pisanty O, Barbosa ICR, Zourelidou M, Regnault T, Crocoll C, Erik Olsen C, Weinstain R, et al. 2016. The Arabidopsis NPF3 protein is a GA transporter. Nature Communications 7: 11486.

Wang J, Kambhampati S, Allen DK, Chen L-Q. 2022a. Comparative metabolic analysis reveals a metabolic switch in mature, hydrated, and germinated pollen in *Arabidopsis thaliana*. Frontiers in Plant Science, in press.

Wang J, Yu Y-C, Li Y, Chen L-Q. 2022b. Hexose transporter SWEET5 confers galactose sensitivity to Arabidopsis pollen germination via a galactokinase. Plant Physiology: kiac068.

Wu L-B, Eom J-S, Isoda R, Li C, Char SN, Luo D, Schepler-Luu V, Nakamura M, Yang B, Frommer WB. 2022. OsSWEET11b, a potential sixth leaf blight susceptibility gene involved in sugar transport-dependent male fertility. New Phytologist, in press.

Xue X, Wang J, Shukla D, Cheung LS, Chen L-Q. 2022. When SWEETs turn tweens: updates and perspectives. Annual Review of Plant Biology 73, in press.

Xue J-S, Yao C, Xu Q-L, Sui C-X, Jia X-L, Hu W-J, Lv Y-L, Feng Y-F, Peng Y-J, Shen S-Y, et al. 2021. Development of the middle layer in the anther of Arabidopsis. Frontiers in Plant Science 12.

Zhang C, Li Y, Wang J, Xue X, Beuchat G, Chen L-Q. 2021. Two evolutionarily duplicated domains individually and post-transcriptionally control SWEET expression for phloem transport. New Phytologist 232: 1793–1807.

Zhang C, Ren M-Y, Han W-J, Zhang Y-F, Huang M-J, Wu S-Y, Huang J, Wang Y, Zhang Z, Yang Z-N. 2022. Slow development allows redundant genes to restore the fertility of *rpg1*, a TGMS line in Arabidopsis. The Plant Journal 109: 1375–1385.

Zheng Y, Deng X, Qu A, Zhang M, Tao Y, Yang L, Liu Y, Xu J, Zhang S. 2018. Regulation of pollen lipid body biogenesis by MAP kinases and downstream WRKY transcription factors in Arabidopsis. PLOS Genetics 14: e1007880.

Zhou X-R, Shrestha P, Yin F, Petrie JR, Singh SP. 2013. AtDGAT2 is a functional acyl-CoA:diacylglycerol acyltransferase and displays different acyl-CoA substrate preferences than AtDGAT1. FEBS Letters 587: 2371–2376.

